# Increased intrinsic membrane excitability is associated with hypertrophic olivary degeneration in spinocerebellar ataxia type 1

**DOI:** 10.1101/2023.10.23.563657

**Authors:** Logan M. Morrison, Haoran Huang, Hillary P. Handler, Min Fu, David D. Bushart, Samuel S. Pappas, Harry T. Orr, Vikram G. Shakkottai

## Abstract

One of the characteristic areas of brainstem degeneration across multiple spinocerebellar ataxias (SCAs) is the inferior olive (IO), a medullary nucleus that plays a key role in motor learning. In addition to its vulnerability in SCAs, the IO is also susceptible to a distinct pathology known as hypertrophic olivary degeneration (HOD). Clinically, HOD has been exclusively observed after lesions in the brainstem disrupt inhibitory afferents to the IO. Here, for the first time, we describe HOD in another context: spinocerebellar ataxia type 1 (SCA1). Using the genetically-precise SCA1 knock-in mouse model (SCA1-KI; both sexes used), we assessed SCA1-associated changes in IO neuron structure and function. Concurrent with degeneration, we found that SCA1-KI IO neurons are hypertrophic, exhibiting early dendrite lengthening and later somatic expansion. Unlike in previous descriptions of HOD, we observed no clear loss of IO inhibitory innervation; nevertheless, patch-clamp recordings from brainstem slices reveal that SCA1-KI IO neurons are hyperexcitable. Rather than synaptic disinhibition, we identify increases in *intrinsic* membrane excitability as the more likely mechanism underlying this novel SCA1 phenotype. Specifically, transcriptome analysis indicates that SCA1-KI IO hyperexcitability is associated with a reduced medullary expression of ion channels responsible for spike afterhyperpolarization (AHP) in IO neurons – a result that has a functional consequence, as SCA1-KI IO neuron spikes exhibit a diminished AHP. These results reveal membrane excitability as a potential link between disparate causes of IO degeneration, suggesting that HOD can result from any cause, intrinsic or extrinsic, that increases excitability of the IO neuron membrane.

**Significance statement:** Little is known about the factors that make inferior olive (IO) neurons susceptible to degeneration in the spinocerebellar ataxias (SCAs), a group of inherited neurodegenerative movement disorders. Another well-described form of IO degeneration, known as hypertrophic olivary degeneration (HOD), results from inhibitory denervation of the IO after brainstem injury. Here, we describe a novel finding of HOD in SCA1 without inhibitory denervation, in association with increased intrinsic membrane excitability and reduced potassium channel transcripts. This suggests that increased membrane excitability may be the underlying primary mechanism of HOD. Identifying hyperexcitability as the mechanistic driver of HOD would imply that reducing intrinsic IO excitability could be an effective strategy for treating diverse causes of both inherited and sporadic olivary degeneration.

## Introduction

The inferior olive (IO) is a nucleus in the medullary brainstem that plays an important role in cerebellar motor learning (Lang et al., 2017). IO neurons send powerful excitatory projections known as climbing fibers into the cerebellum, where they synapse onto Purkinje cell dendrites to generate cerebellar complex spikes (De Zeeuw et al., 1998; Ausim Azizi, 2007; Ito, 2013; Llinas, 2013). The IO is susceptible to degeneration in a number of conditions, including sporadic olivopontocerebellar atrophy (OPCA) (Konigsmark and Weiner, 1970; Duvoisin, 1984) and the spinocerebellar ataxias (Seidel et al., 2012). In addition, IO neurons are vulnerable to a pathology known as hypertrophic olivary degeneration (HOD). Clinically, HOD occurs secondary to lesions in the brainstem (pontine hemorrhage, most commonly (Smets et al., 2017)) that disrupt inhibitory fibers that travel from the contralateral dentate nucleus of the cerebellum and through the brainstem to synapse onto the IO (a white matter tract known as the Guillain-Mollaret triangle (Ogut et al., 2023)). An acute lesion to these fibers results in an initial increase in IO neuron soma size, generally observed as an increase in gross IO volume (Jellinger, 1973; Ruigrok et al., 1990; Wang et al., 2019). This is ultimately followed by cell loss, gliosis, and atrophy of the IO, a process that sometimes extends for years after the initial insult (Goto and Kaneko, 1981; Pandey et al., 2013). Although HOD ultimately results in IO degeneration, its pathology is considered distinct from the IO atrophy observed in OPCA and the spinocerebellar ataxias due to the lack of any clinically-evident hypertrophy in these disorders (Koeppen, 2018).

The spinocerebellar ataxias (SCAs) are a group of inherited neurodegenerative diseases that cause a progressive loss of motor function (Paulson, 2009; Seidel et al., 2012). Progressive brainstem degeneration (including in the IO) is known to occur in SCA1, SCA2, SCA3, and SCA7, which together account for the majority of SCA cases (Rub et al., 2013). This pathology manifests late in disease progression, corresponding closely to the onset of breathing and swallowing deficits (symptoms that can eventually cause premature death, especially in SCA1) (Durr, 2010). Among these common SCAs, the fastest-progressing is SCA1 (Scott et al., 2020), a subtype that exhibits IO degeneration as a characteristic pathology (Koeppen et al., 2013). The mechanisms that underlie this degeneration, as well as brainstem pathology in the SCAs broadly, remain poorly understood.

In this study, we examined IO morphology and physiology in the genetically-precise SCA1 knock-in (SCA1-KI) mouse model. For the first time, we have identified changes consistent with early HOD in a model not of brainstem injury, but of neurodegenerative disease. In neurons of the principal olivary nucleus (IOPr) of SCA1-KI mice, we observed early dendritic hypertrophy followed by late somatic hypertrophy. We found that these morphological changes are concurrent with a loss of immunoreactivity for the calcium-binding protein calbindin. Calbindin loss is characteristic of SCA1 pathology in both IO neurons (Koeppen et al., 2013; Yu et al., 2014) and Purkinje cells (Vig et al., 1998; Koeppen, 2005) and has been used as a surrogate for neurodegeneration in various mouse models of the disease, including SCA1-KI mice (Burright et al., 1995; Watase et al., 2002). Interestingly, structural and functional analyses of IO innervation demonstrate no loss of inhibitory input, suggesting an intrinsic mechanism of HOD in SCA1 that is distinct from the extrinsic mechanism produced by lesions of the brainstem. Using patch-clamp electrophysiology and unbiased transcriptome analysis, we further found that SCA1-KI IO neurons are hyperexcitable, likely due to a reduced expression of ion channel genes – specifically, genes that encode certain calcium and potassium channels that are known to regulate IO neuron intrinsic excitability by mediating spike afterhyperpolarization (AHP) (Llinas and Yarom, 1981b, a). Together, these results reveal intrinsic membrane excitability as a potential link between disparate causes of HOD, suggesting that hyperexcitability from any cause, extrinsic or intrinsic, converges on the same mechanistic pathway of IO degeneration.

## Materials and Methods

### Mouse studies

All animal studies were reviewed and approved by the Institutional Animal Care and Use Committee (IACUC) at the institution where they were performed (University of Michigan, University of Texas Southwestern Medical Center, or University of Minnesota) and were conducted in accordance with the United States Public Health Service’s Policy on Humane Care and Use of Laboratory Animals. SCA1 knock-in (SCA1-KI) mice (RRID:MGI:3774931), which express an expanded CAG triplet repeat in the endogenous *Atxn1* locus (Watase et al., 2002), were maintained on a C57BL/6 background. SCA1-KI mice were heterozygous for the expanded *Atxn1* allele (*Atxn1^154Q/2Q^*), with wild-type littermates (*Atxn1^2Q/2Q^*) used as controls. SCA1 transgenic (SCA1-Tg) mice (RRID:MGI:5518618) overexpress the human *ATXN1* gene with an expanded CAG triplet repeat under the Purkinje cell-specific murine *Pcp2 (L7)* promotesr (Burright et al., 1995) and were maintained on an FVB background (Jackson Labs, RRID:IMSR_JAX:001800). SCA1-Tg mice were homozygous for the transgene (*ATXN1[82Q]^tg/tg^*), with age/sex-matched wild-type FVB mice used as controls. For both mouse models, studies were performed at either 13-15 weeks of age (defined as the “14 week” timepoint) or 29-32 weeks of age (defined as the “30 week” timepoint). Sexes were balanced for all animal studies.

### Patch-clamp electrophysiology

#### Solutions

Artificial cerebrospinal fluid (aCSF) used in these experiments contained: 125 mM NaCl, 3.8 mM KCl, 26 mM NaHCO_3_, 1.25 mM NaH_2_PO_4_, 2 mM CaCl_2_, 1mM MgCl_2_, and 10 mM glucose. For all recordings, pipettes were filled with internal solution containing: 140 mM K-Gluconate, 4 mM NaCl, 10^-3^ mM CaCl_2_, 4mM Mg-ATP, 10^-2^ mM EGTA, 10 mM HEPES, at pH 7.3 and osmolarity ∼285 mOsm, as described previously for IO recordings (Lefler et al., 2014).

#### Preparation of brain slices for acute electrophysiological recordings

Slices were prepared using the “hot cut” technique (Huang and Uusisaari, 2013). This process, which involves tissue sectioning in standard aCSF at physiological temperatures rather than a sucrose-rich aCSF at near-freezing temperatures, is a recent advancement that has allowed for patch-clamp recordings in brain regions previously deemed inaccessible in adult mice due to their hypersensitivity to slicing-induced damage. This technique has facilitated studies of nuclei never-before investigated in the adult mouse brain, including the inferior olive (Lefler et al., 2014). Briefly, mice were deeply anesthetized by isoflurane inhalation, decapitated, and brains were rapidly removed and submerged in pre-warmed (33°C) aCSF. 300 μm coronal slices were prepared on a VT1200 vibratome (Leica Biosystems, Deer Park, IL) in aCSF held at 32.5°C-34.0°C during sectioning. Slices were then incubated in carbogen-bubbled (95% O2, 5% CO2) aCSF at 33°C for 45 min. Slices were then stored in carbogen-bubbled aCSF at room temperature (RT) until use. During recording, slices were placed in a recording chamber and continuously perfused with carbogen-bubbled aCSF at 33°C at a flow rate of 2.5 mL/min.

#### Patch-clamp recordings

IO neurons were visually identified for patch-clamp recordings using a 40x water immersion objective and a Nikon Eclipse FN1 upright microscope with infrared differential interference contrast (IR-DIC) optics. Identified cells were visualized using NIS Elements image analysis software (Nikon, Melville, NY). Patch pipettes were pulled to resistances of ∼3 MΩ from thin-walled glass capillaries (World Precision Instruments, Cat# 1B120F-4). Importantly, we found that pipette shape had an outsized impact on the success of these difficult recordings; as such, special care was taken to make pipettes with a short taper and a perfectly flat, ∼1 μm diameter tip. Data were acquired using a CV-7B headstage amplifier, a Multiclamp 700B amplifier, a Digidata 1440A interface (Axon Instruments, San Jose, CA), and pClamp-10 software (Molecular Devices, San Jose, CA). All data were digitized at 100 kHz. All voltages were corrected for liquid gap junction potential, previously calculated to be 10 mV (Dell’Orco et al., 2015).

### Purkinje cell *in-vivo* electrophysiology

#### Surgical preparation of craniotomy site

*In-vivo* electrophysiological recordings were performed with custom equipment and techniques as described previously (Heiney et al., 2014). Briefly: 14-week-old wild-type and SCA1-KI mice were surgically prepared for recordings one day in advance. Isoflurane anesthesia (dosage 4% for induction, 1.5% for maintenance) was used during all surgical procedures. After a scalp incision, two 0.6 x 0.06” SL flat SS anchor screws (Antrin, Fallbrook, CA) were placed into the skull slightly posterior to bregma. A custom titanium headplate was then affixed to both the skull and the anchor screws with C&B-Metabond (Parkell, Edgewood, NY), and a recording chamber was shaped around the skull above the cerebellum using dental cement (Lang Dental, Wheeling, IL). Two 3 mm craniotomies (centered at −6.2 AP, ±2.1 ML from Bregma) were performed to expose the anterior lobules of each cerebellar hemisphere. Craniotomy sites were protected with Kwik-Sil (World Precision Instruments, Sarasota, FL) until recording.

#### Recording procedure

On the day of recording, animals were secured to the recording platform using the titanium headplate described above. All recordings were performed in awake mice in a single session per mouse, with the total duration of the session limited to 1 hr. After the headplate was attached to the platform, the dental cement recording chamber was filled with sterile PBS and a 2.5-3.5 MΩ tungsten electrode (Thomas Recording, Giessen, Germany) was inserted into one of the craniotomy sites to acquire data. The recording electrode was slowly lowered into the cerebellar cortex until Purkinje cells were identified by their characteristic physiology (20-80 Hz simple spikes and the presence of complex spikes). From this point, the electrode was lowered in 10 μm increments until maximum simple spike amplitude was reached. Though electrode placement varied between animals, final electrode depth was typically 2-3 mm from the surface of the brain. Purkinje cell activity was recorded using a DP-301 differential amplifier (Warner Instruments, Holliston, MA), digitized at 100 kHz using a Digidata 1440A interface (Axon Instruments, San Jose, CA), and analyzed using pClamp-10 software (Molecular Devices, San Jose, CA). A minimum of 5 min of activity was recorded per craniotomy site. Recording position in the cerebellar cortex was verified by assessing tissue damage from the recording electrode via Nissl staining.

#### Data analysis

Complex spike frequency was analyzed in 10 Hz high-pass filtered traces, with the criterion for identification that complex spikes must exhibit a voltage higher than the maximum simple spike peak for the entire trace, as described previously (Warnaar et al., 2015). Displayed traces from *in-vivo* electrophysiological recordings were first filtered to reduce minor 60 Hz electrical interference, then high-pass filtered at 10 Hz.

### Cell filling and morphological analysis

#### Filling protocol and section preparation

Before patch clamp recordings, Alexa Fluor 488 Succinimidyl Ester (Invitrogen, Cat# A20000) powder was dissolved separately in internal solution at a concentration of 1 mg/mL. Pipette tips were filled with a miniscule amount of standard internal solution so fluorescent dye would not be washed over the tissue when attempting to patch. This was accomplished by placing pipettes upside down (i.e., tip up) in a microcentrifuge tube containing ∼20 μL internal solution for ∼5 min, which filled the pipette tip by capillary action. From here, pipettes were backfilled with the fluorescent internal solution, allowing cells to be filled with dye while recording. All cells were filled for 15-20 min and confirmed as filled after pipette removal by the presence of intense somatic fluorescence at 40X (as shown in **Fig. 5A, inset**). Only one cell per side was filled on each section to avoid tracing errors from dendrite overlap. In addition, a scalpel was used to make a small notch in the left spinal trigeminal tract of each section to mark its orientation. After filling was complete, free-floating brainstem sections were immediately fixed in 1% PFA for 1 hr at RT, then washed 3x in PBS. To avoid crushing the sections during mounting (as patch-clamp slices are much thicker than typical histological sections) a square was drawn on the slide using clear nail polish and allowed to dry immediately before attempting to mount the tissue. Sections were mounted in this square using ProLong Gold Antifade Mountant with DAPI (Cell Signaling Technology, Cat# 8961S), with specific attention being paid to the notch in the section to ensure that the surface of the section with the filled cell was mounted against the coverslip, not the slide.

#### Confocal imaging and of filled cells

Sections were imaged using a Nikon C2+ microscope with a 63x oil immersion objective. Imaging of each cell was accomplished by taking multiple z-stack images at the microscope’s minimum of 0.207 μm between each image of the stack.

#### 3D reconstruction of cells and morphological analysis

After imaging, 3D reconstructions of cells were generated from z-stack images using the open-source tracing software Vaa3D (Peng et al., 2010; Peng et al., 2014a; Peng et al., 2014b). For each cell, a conservative initial trace was generated automatically by Vaa3D, then manually reviewed and appended to include all visible projections of the neuron. All single-cell morphological calculations were made automatically by Vaa3D, with dendritic tip and bifurcation counts confirmed as accurate by manual review.

### Immunohistochemistry

#### Sample preparation and imaging

Mice were anesthetized under isoflurane inhalation and brains were removed, fixed in 1% paraformaldehyde (PFA) for 1 hour, after which they were placed in 30% sucrose in phosphate-buffered saline (PBS) for 48 hours at 4°C. Brains were then cryo-embedded in a 1:1 mixture of 30% sucrose (in PBS) and OCT compound (Fisher Scientific, Cat# 23-730-571) at −80°C for at least 24 hrs. Tissue blocks were sectioned to 20 µm thickness on a CM1850 cryostat (Leica Biosystems, Deer Park, IL). Tissue was permeabilized with 0.4% triton in PBS, then non-specific binding was minimized by blocking with 5% normal goat serum in 0.1% triton in PBS. Primary antibodies (Abs) were incubated in PBS containing 0.1% triton and 2% normal goat serum at 4°C overnight, while secondary Abs were applied for one hour at room temperature in PBS.

#### Antibodies used

To stain for neuronal nuclear protein (NeuN) for stereology (**Figs. 2 & 4**), Rabbit anti-NeuN primary Ab (1:500, Cell Signaling, Cat# 12943, RRID:AB_2630395) and Goat anti-Rabbit IgG (H+L) Alexa Fluor 488-conjugated secondary Ab (1:200; Invitrogen, Cat# A11034, RRID:AB_2576217) were used. To stain for Calbindin (Calb) for stereology (**Figs. 2 & 4**), Mouse anti-Calb primary Ab (1:1000; Sigma Aldrich, Cat# C9848, RRID:AB_476894) and Goat anti-Mouse IgG (H+L) Alexa Fluor 594-conjugated secondary Ab (1:200; Invitrogen, Cat# A11005, RRID:AB_2534073) were used. To stain for glutamate decarboxylase (GAD) for assessment of IO inhibitory innervation (**Figs. 3 & 4**), Rabbit anti-GAD primary Ab (1:500, Sigma-Aldrich, Cat# G5163, RRID:AB_477019) and Goat anti-Rabbit IgG (H+L) Alexa Fluor 594-conjugated secondary Ab (1:200, Invitrogen, Cat# A11012, RRID:AB_2534079) were used. To stain for Calbindin (Calb) for assessment of IO inhibitory terminals (**Figs. 3 & 4**), Mouse anti-Calb primary Ab (1:1000; Sigma Aldrich, Cat# C9848, RRID:AB_476894) and Goat anti-Mouse IgG (H+L) Alexa Fluor 488-conjugated secondary Ab (1:200; Invitrogen, Cat# A11001, RRID:AB_2534069) were used. To stain for small conductance calcium-activated potassium channel 2 (SK2) in IO ion channel studies (**Fig. 7**), Rat anti-SK2 primary Ab (1:200, NeuroMab clone K78/29, Cat# 75-403, RRID:AB_2877597) and Goat anti-Rat IgG (H+L) Alexa Fluor 488-conjugated secondary Ab (1:200, Invitrogen, Cat# A11006, RRID:AB_2534074) were used. To stain for Calbindin (Calb) for IO ion channel studies (**Fig. 7**), Mouse anti-Calb primary Ab (1:1000; Sigma Aldrich, Cat# C9848, RRID:AB_476894) and Goat anti-Mouse IgG (H+L) Alexa Fluor 594-conjugated secondary Ab (1:200; Invitrogen, Cat# A11005, RRID:AB_2534073) were used.

#### Fluorescence intensity measurements

To measure the intensity of GAD staining (**Figs. 3 & 4**), images acquired at 10x magnification were used. Fluorescence intensity analysis was performed using ImageJ. A rectangular box was placed in the IO, specifically in the area of the principal olivary nucleus (IOPr). Mean pixel intensity was measured for each rectangle, which was then used as the raw fluorescence intensity value for each section. Box size was identical in all cases and placed in similar areas of the IOPr between sections. Two sections were imaged per animal and the mean of their fluorescence values were used as the fluorescence intensity for that animal. All tissue processing and imaging was performed at the same session and microscope settings were identical for all acquired images. During imaging and analysis, the experimenter was blind to genotype.

#### Confocal microscopy

Imaging was performed on a Nikon C2+ confocal microscope. Single-plane images were acquired at 63x magnification using an oil-immersion lens, with microscope settings kept constant between all samples under each set of antibodies. Samples were prepared and imaged with the experimenter blind to genotype.

### Inferior olive stereology

#### Cell counts

Estimates for the total number of neurons (NeuN^+^ cells) in the IO, total number of healthy neurons (NeuN^+^,Calb^+^ cells) in the IO, and total IO volume were quantified with an unbiased stereological approach using the optical fractionator probe in Stereoinvestigator (MBF Bioscience, Williston, VT). 30 μm serial coronal sections through the medulla were mounted on slides and co-stained for NeuN and Calbindin as described above. Sections were observed under epifluorescence on a Zeiss Axioimager M2 microscope, where regions of interest were first outlined using a 10x objective lens. 10 sections per brain were analyzed, with a section evaluation interval of 4. Cells within the outlined region were counted using a 63x oil immersion objective, with a ∼20 μm counting depth and 1 μm guard zones. A 200 μm x 200 μm counting frame and 50 μm x 50 μm sampling grid were used for all counts, determined effective in pilot studies such that the Gunderson coefficient of error was less than 0.1 for all markers. The top of each stained cell body was the point of reference. The pyramids (py), ventral gigantocellular reticular nucleus (GRN), lateral paragigantocellular nucleus (PGRNl), medial lemniscus (ml), and magnocellular reticular nucleus (MARN) were used as anatomical boundaries for the IO.

#### Cell size measurements

IOPr neuron cell size was quantified by measuring the average soma area (μm^2^) for 30-50 cells per animal. Measurements were made concurrent with stereological analysis using the 63x oil immersion objective and the 4-ray nucleator probe in Stereoinvestigator (MBF Bioscience, Williston, VT). Cell size was measured while visualizing cells with NeuN. Analysis was constrained to the principal nucleus of the inferior olive (IOPr), identified by its distinct laminar structure. 200–400 neurons (from 5–8 brains) were measured for each genotype at each timepoint.

### Transcriptome analysis

#### Administration of antisense oligonucleotides (ASOs)

ASO treatment was performed in SCA1-KI and wild-type mice at 5 weeks of age by intracerebroventricular (ICV) injection. For these injections, mice were anesthetized with an intraperitoneal (IP) ketamine/xylazine cocktail (100 mg/kg ketamine and 10 mg/kg xylazine). Using a stereotax, the cranium was burr drilled and a Hamilton Neuros Syringe (65460-05) was positioned at the following coordinates: AP, 0.3; ML, 1.0; DV, –2.7 mm from bregma. Injections of 10 μl ASO 618353 (50 μg/μl) dissolved in PBS without Ca/Mg (Gibco, 14190) or vehicle were delivered via a micro-syringe pump at 25 nl/s. Immediate postoperative care included subcutaneous delivery of 250 μl saline, carprofen (7 mg/kg) and Buprenorphine SR (Zoopharm Pharmacy) (2 mg/kg).

#### RNA isolation and sequencing

Total RNA was isolated from dissected medullae of 28-week-old wild-type vehicle-injected mice (8 samples), 28-week-old SCA1-KI vehicle injected mice (8 samples), and 28-week-old SCA1-KI ASO-injected mice (7 samples) using TRIzol Reagent (Life Technologies) following the manufacturer’s protocols. Tissue was homogenized using RNase-Free Disposable Pellet Pestles (Fisher Scientific) in a motorized chuck. For RNA-seq, RNA was further purified to remove any organic carryover using the RNeasy Mini Kit (Qiagen) following the manufacturer’s RNA Cleanup protocol. Purified RNA was sent to the University of Minnesota Genomics Center for quality control, including quantification using fluorimetry (RiboGreen assay, Life Technologies), and RNA integrity was assessed with capillary electrophoresis (Agilent BioAnalyzer 2100, Agilent Technologies Inc.), generating an RNA integrity number (RIN). All submitted samples had greater than 1 μg total mass and RINs greater than or equal to 7.9. Library creation was completed using oligo-dT purification of polyadenylated RNA, which was reverse transcribed to create cDNA. cDNA was fragmented, blunt ended, and ligated to barcoded adaptors. Library was size-selected to 320 bp ±5% to produce average inserts of approximately 200 bp, with size distribution validated using capillary electrophoresis and quantified using fluorimetry (PicoGreen, Life Technologies) and qPCR. Libraries were then normalized, pooled, and sequenced on an Illumina HiSeq 2000 using a 100-nt paired-end read strategy. Data were stored and maintained on University of Minnesota Supercomputing Institute Servers. Reads were aligned to the mouse reference genome (GRcm38) with hisat2 (Kim et al., 2015) using default parameters for stranded libraries, the hisat2 GRCm38 index for the genome plus SNPs, and transcripts and using Ensembl’s release 87 of the GRCm38 gene annotations. Read counts per gene were summed using featureCounts from the subread package (Liao et al., 2014). Genes less than about 300 bp are too small to be accurately captured in standard RNA-seq library preparations, so they were discarded from downstream analyses. Differential expression analysis was carried out using the R package DESeq2 (Love et al., 2014). Medulla samples were collected and sequenced in two batches, with batch effect corrected for in the differential expression analysis model. Genes with a false discovery rate (FDR) value ≤ 0.05 were considered significant. For both data sets, genes were ranked by –log_10_(*P* value) using differential expression comparison between the wild-type vehicle-injected and SCA1-KI ASO–injected groups, with the sign of the fold change assigned to the ranking metric; i.e., if the fold change was negative, the ranking metric was –1 × (–log_10_[*P* value]). Statistically significant differentially-expressed genes (DEGs) were identified as all genes with *P* < 0.05 (or *P_adjusted_* < 0.05, where appropriate) and an expression value of either log_2_FoldChange < −0.3 or log_2_FoldChange > 0.3.

### Experimental Design and Statistical Analysis

Experimenters were kept blind to genotype for all experiments to avoid biasing results. To prevent variability within experiments, every individual analysis was performed by the same experimenter (e.g., all GAD staining analysis was performed by the same person) and using the same conditions/settings (e.g., all staining for an experiment was done in a single session and analyzed with the same microscope settings) for that experiment’s full mouse cohort. Individual statistical tests are described in the figure legends for all data. As this was an exploratory study, the null hypotheses were not prespecified and calculations for statistical power were not performed prior to study initiation. It follows from the exploratory nature of the experiments that calculated *P* values cannot be interpreted as hypothesis testing but only as descriptive. Though formal power analysis was not performed, we estimated sample size based on our (and others’) previous work in SCA1 mouse models. Number of individual datapoints and total number of animals for all experiments are reported in figure legends. Statistical analysis was done using Microsoft Excel, Prism 6.0 (GraphPad Software, Boston, MA), and SigmaPlot (Systat Software, Palo Alto, CA), with statistical significance defined as *P* < 0.05. If statistical significance was achieved, we performed post-hoc analysis corresponding to the experiment, as specified, to account for multiple comparisons. All *t*-tests were two-tailed Student’s *t*-tests, with the level of significance (alpha) set at 0.05. Enrichment of ion channels was calculated using Fisher’s exact test, with initial enrichment calculated by comparing ion channel gene proportion amongst DEGs to ion channel gene proportion amongst all genes analyzed (270 potential mouse ion channel transcripts of 30,973 total transcripts assessed) in a 2 x 2 contingency table.

## Results

### Neurons of the principal olivary nucleus in SCA1 mice undergo degenerative hypertrophy

To assess the morphological changes of neurons in the principal olivary nucleus (IOPr) due to SCA1, we filled individual IOPr neurons from coronal brainstem slices of 14-week-old SCA1 knock-in (SCA1-KI) mice. This mouse strain was used because it is the most genetically precise model of SCA1, generated by the insertion of 154 CAG repeats into the endogenous *Atxn1* locus. This causes slowly-progressing movement phenotypes that mirror human SCA1: SCA1-KI motor incoordination begins at 5 weeks (Watase et al., 2002), becomes robust by 14 weeks (Bushart et al., 2021), and is severe by 30 weeks (Cvetanovic et al., 2011). Importantly, these mice also model brainstem phenotypes (unlike previous SCA1 mouse models), as they produce mutant ATXN1 in all cell types in which *Atxn1* is endogenously expressed. We filled cells from both SCA1-KI and wild-type littermate controls with a biologically inert fluorescent dye, then imaged them in three dimensions using z-stack confocal microscopy. Using the open-source imaging software suite Vaa3D (Peng et al., 2010; Peng et al., 2014a; Peng et al., 2014b), we generated 3D reconstructions from these z-stacks to measure various parameters of each cell’s shape and size.

Neurons of the IOPr are multipolar, generally described as having a “cloud” of dendrites surrounding the soma in three dimensions (De Zeeuw et al., 1998; Vrieler et al., 2019). At 14 weeks, we found that these IOPr dendritic arbors are significantly larger and more complex in SCA1-KI mice than in wild-type controls (**Fig. 1A**). SCA1-KI IOPr neurons have a greater number of dendritic tips and bifurcations (**Fig. 1B-C**), a larger total length of dendrites (**Fig. 1D**), and a significantly higher fractal dimensionality (a measure of the complexity of a three-dimensional object) (**Fig. 1E**). Though this indicates significant hypertrophy of SCA1-IOPr neuron dendrites, we did not observe a change in the total span of the dendritic arbor (**Fig. 1F-H**). Similarly, soma surface area was largely unchanged between genotypes (**Fig. 1I**).

**Figure 1.**
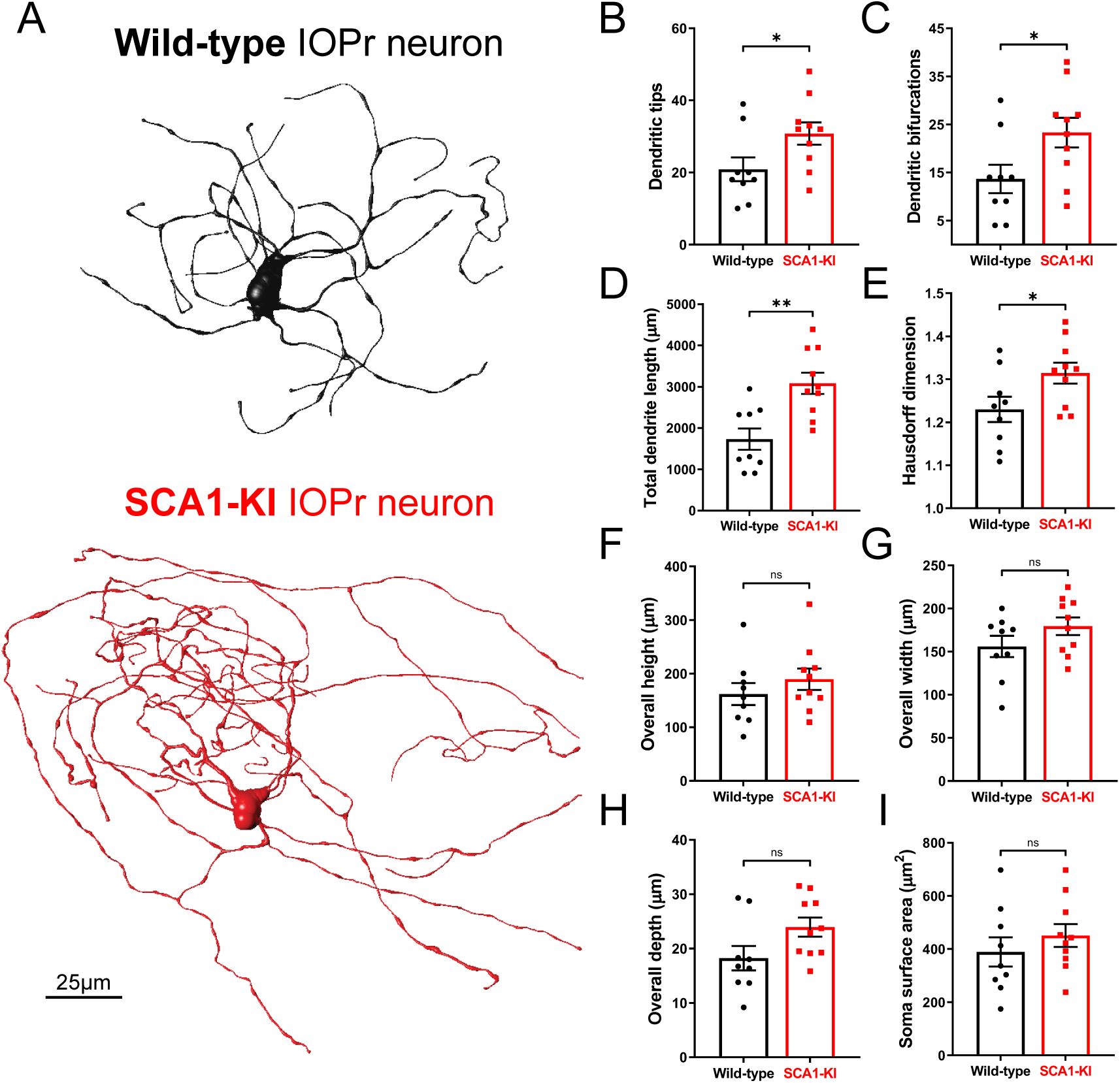
IOPr neuron dendritic arbors are hypertrophic in SCA1-KI mice. **(A)** Representative morphological reconstructions of dye-filled IO neurons in wild-type (top) and SCA1-KI (bottom) mice at 14 weeks of age (an early symptomatic timepoint). **(B-I)** The dendritic arbors of SCA1-KI IO neurons are larger and more complex than wild-type controls. SCA1-KI IO neurons exhibit a greater number of dendritic tips **(B)** and dendritic bifurcations **(C)**, as well as a marked increase in total dendritic length **(D)** and Hausdorff dimension **(E)** (a three-dimensional measure of object complexity). The overall span of IO neurons, shown as height **(F)**, width **(G)**, and depth **(H)**, is largely unchanged between the two genotypes. No change in IO soma size (measured by surface area) was detected between the two genotypes **(I)**. Data are expressed as mean ± SEM. Statistical significance derived by unpaired *t*-test with Welch’s correction, * = *P* < 0.05, ** = *P* < 0.01, ns = not significant. *N_wild-type_* = 9 cells from 7 mice, *N_SCA1-KI_* = 10 cells from 7 mice.

To investigate SCA1-related phenotypes in the IO as a whole, we quantified IO cell number in 14-week-old (early symptomatic) and 30-week-old (late symptomatic) mice using unbiased stereology. This was performed using brainstem sections co-stained for neuronal nuclear protein (NeuN; a marker of all neurons) and calbindin (Calb). Calb is a well-conserved calcium-binding protein that plays a critical role in calcium buffering in Purkinje cells (Schwaller et al., 2002). Historically, loss of Calb-immunoreactive (Calb^+^) Purkinje cells has been widely used as a surrogate for Purkinje cell degeneration in SCA mouse models, including models of SCA1 (Watase et al., 2002), SCA2 (Hansen et al., 2013), SCA3 (Switonski et al., 2015), and SCA17 (Cui et al., 2017). Mouse IO neurons also exhibit high levels of Calb expression (Yu et al., 2014), and loss of Calb^+^ cells in the IO has similarly been used as an indicator of degeneration in the IO of both ataxic mice (Zanjani et al., 2004) and human SCA patients (Koeppen et al., 2013). Here, we found that 14-week-old SCA1-KI mice exhibit a ∼25% loss of Calb^+^ IO neurons (**Fig. 2E**) without a discernible decrease in total cell number (**Fig. 2D**) or mean IO volume (**Fig. 2F**), suggesting the presence of early IO neurodegeneration in these mice. 30-week-old animals also showed no difference in total IO cell number (**Fig. 2K**) or mean IO volume (**Fig. 2M**), though loss of Calb^+^ neurons in the SCA1-KI IO was not significant (*P* = 0.0658) at this stage (**Fig. 2L**).

**Figure 2.**
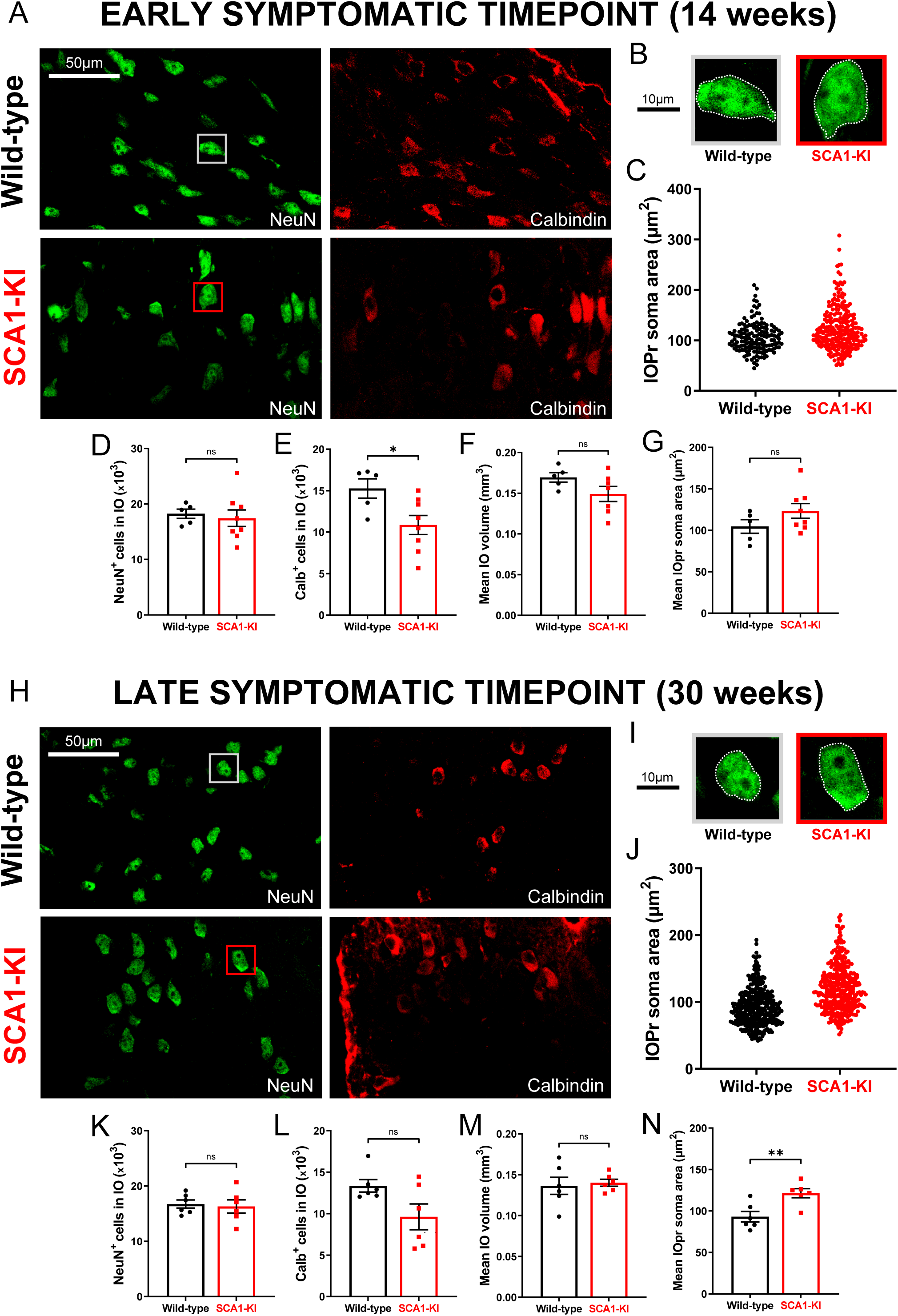
IO neurons in SCA1-KI mice exhibit somatic hypertrophy and degeneration. **(A)** Coronal histological sections showing IOPr neurons in wild-type (top) and SCA1-KI (bottom) mice at an early symptomatic timepoint of 14 weeks. Sections have been stained for NeuN (green) and Calbindin (red). **(B)** Inset showing representative IOPr cells of each genotype. **(C, G)** IOPr neuron soma size was estimated by measuring the area of the soma in ∼200 cells per genotype. No change in IOPr soma size in SCA1-KI mice is evident at the 14 week timepoint. **(D-F)** Quantification of unbiased stereological analysis, providing an estimate of both total cell number and gross volume for the entire IO. At 14 weeks, degeneration is already apparent in the SCA1-KI IO, shown by loss of calbindin signal. **(H)** Coronal histological sections showing principal IO in wild-type (top) and SCA1-KI (bottom) mice at a late symptomatic timepoint of 30 weeks. Sections have been stained for NeuN (green) and Calbindin (red). **(I)** Inset showing representative IOPr cells of each genotype. **(J, N)** IOPr neuron soma size was estimated by measuring the area of the soma in ∼400 cells per genotype. At the 30 week timepoint, SCA1-KI IOPr neurons exhibit a highly significant increase in soma size **(J)**, with a shift in the entire distribution of neurons towards a larger size **(N)**. **(K-M)** Quantification of unbiased stereological analysis reveals an even greater loss of calbindin signal at 30 weeks **(K-M)**, indicating a more severe degenerative state. Data are expressed as mean ± SEM. Statistical significance derived by unpaired *t*-test with Welch’s correction, * = *P* < 0.05, ** = *P* < 0.01, ns = not significant. 14 weeks: *N_wild-type_* = 5 mice, *N_SCA1-KI_* = 8 mice; 30 weeks: *N_wild-type_* = 6 mice, *N_SCA1-KI_* = 6 mice.

While performing stereological quantifications on these sections, we also conducted cell size measurements. Focusing solely on the IOPr, we recorded the soma area of 200-400 cells per genotype at each timepoint (**Fig. 2C,J**). At 14 weeks, we found no difference in the average soma area of IOPr neurons between genotypes (**Fig. 2G**), a result that is consistent with soma area measurements from filled cells at 14 weeks (**Fig. 1I**). However, at 30 weeks, IOPr soma areas were ∼25% larger on average in SCA1-KI mice compared to wild-types (**Fig. 2N**), indicating significant somatic hypertrophy. These results, along with the morphological changes observed in single IOPr neurons at 14 weeks (**Fig. 1**), suggest an SCA1-associated hypertrophy in IOPr neurons that appears first in dendrites and later in the soma.

This concurrence of hypertrophy and degeneration is uncommon amongst neuronal disorders. Interestingly, however, the IO is one of the few brain regions in which degenerative hypertrophy has been previously described. This phenomenon, known as hypertrophic olivary degeneration (HOD), can occur if brainstem injury (usually pontine hemorrhage (Smets et al., 2017)) causes substantive loss of inhibitory input to the IO (Wang et al., 2019). The degenerative hypertrophy identified here appears strikingly similar to this well-described pathology, suggesting that SCA1-associated IO dysfunction may constitute a novel cause of HOD.

### Degenerative hypertrophy in SCA1-KI IOPr neurons is cell-autonomous

In order to rule out other potential causes of HOD, we analyzed inhibitory projections to the IOPr in SCA1-KI and wild-type mice at 14 weeks and 30 weeks. Coronal brainstem sections were stained for glutamate decarboxylase (GAD), a marker of GABAergic terminals (**Fig. 3A,C**). At both 14 weeks and 30 weeks, there was no change in the intensity of GAD staining between groups (**Fig. 3B,D**), indicating that inhibitory input to the IOPr remains structurally intact during IO degeneration. To assess whether these inputs are functionally intact, we performed *in-vivo* recordings of cerebellar Purkinje cells (PCs) in awake, head-fixed SCA1-KI and wild-type mice at 14 weeks. The primary output of IO neurons is the complex spike (CS), a characteristic high-amplitude depolarization of the PC membrane that occurs when climbing fibers are activated (Davie et al., 2008; Streng et al., 2018). These CSs can be observed *in-vivo* as distinctly large depolarizing membrane deflections, as shown in our recordings (**Fig. 3E**). During these recordings, we observed significantly fewer CSs in SCA1-KI PCs compared to wild-type PCs (**Fig. 3F**). This suggests that SCA1-KI IOPr neurons are indeed not experiencing a loss of inhibitory tone, as we would expect disinhibition of the IO to cause an increase in final IO output onto PCs (as previously described (Lang et al., 1996)).

**Figure 3.**
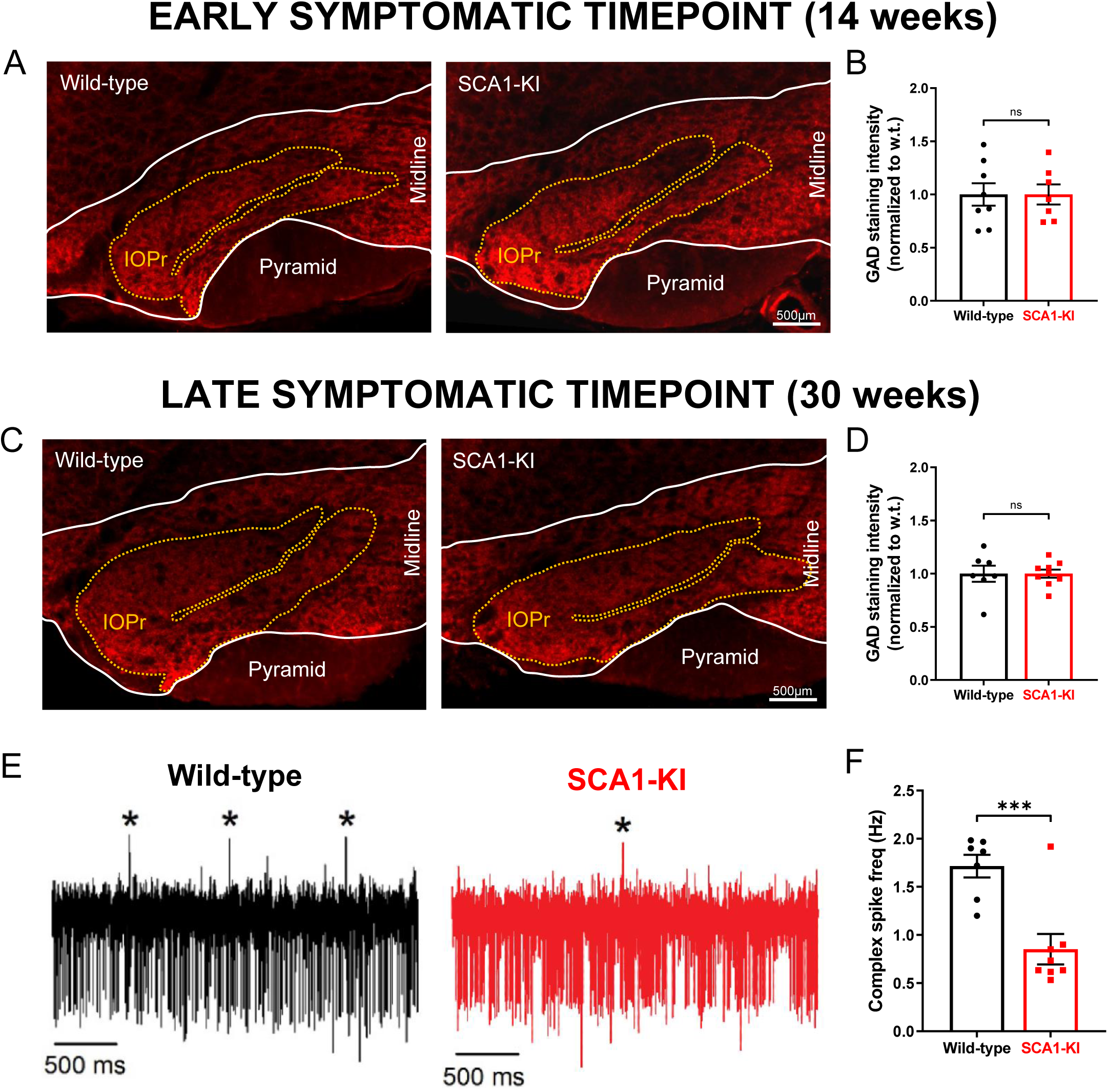
Inhibitory synaptic inputs to the IO remain intact in SCA1-KI mice. **(A)** Coronal histological sections showing the IOPr in wild-type (left) and SCA1-KI (right) mice at an early symptomatic timepoint of 14 weeks. Sections have been stained for glutamate decarboxylase (GAD), a marker of GABAergic terminals (red). **(B)** Quantification of staining reveals that GAD signal is retained in the SCA1-KI IO at 14 weeks. **(C)** Coronal histological sections showing GAD staining in the principal IO at a late symptomatic timepoint of 30 weeks. **(D)** Quantification of staining reveals that GAD signal is also retained in the SCA1-KI IO at 30 weeks. **(E)** Representative traces of *in-vivo* cerebellar spiking patterns in head-fixed, awake mice at 14 weeks. Complex spikes (generated by IO neurons) are indicated with asterisks (*). **(F)** Complex spike frequency is reduced in SCA1-KI cerebella, suggesting that IO neurons *in-vivo* are not disinhibited by loss of inhibitory synaptic input. Data are expressed as mean ± SEM. Statistical significance derived by unpaired *t*-test with Welch’s correction, *** = *P* < 0.001, ns = not significant. **(A-D)** 14 weeks: *N_wild-type_* = 8 mice, *N_SCA1-KI_* = 7 mice; 30 weeks: *N_wild-type_* = 7 mice, *N_SCA1-KI_* = 9 mice. **(E,F)** *N_wild-type_* = 7 mice, *N_SCA1-KI_* = 8 mice.

Because SCA1-KI mice express mutant *Atxn1* via its endogenous promoter, ATXN1 function is altered broadly across the SCA1-KI mouse brain. Thus, it is possible that the observed IO degenerative hypertrophy is being influenced by other affected brain regions within the IO circuit. To determine whether *Atxn1* dysfunction outside of the IO is contributing to this degenerative hypertrophy, we assessed cell number, cell size, and the density of inhibitory inputs in SCA1 transgenic mice (SCA1-Tg). These mice overexpress the human *ATXN1* gene with an expanded CAG triplet repeat under the murine *Pcp2 (L7)* promoter (Burright et al., 1995). This drives selective expression of polyglutamine-expanded ATXN1 (82 repeats) in cerebellar Purkinje cells, which results in a more severe motor phenotype than observed in SCA1-KI mice. This includes an earlier age-of-onset of motor symptoms, such that the 14 week timepoint is already late in the symptomatic progression of SCA1-Tg mice (Clark et al., 1997). Unbiased stereological quantification reveals that SCA1-Tg mice do not show signs of IO degeneration at 14 weeks, demonstrated by no loss of Calb^+^ cells (**Fig. 4E**). Interestingly, though, mean IO volume (**Fig. 4F**) and mean NeuN^+^ cell count (**Fig. 4D**) both exhibited a significant increase in SCA1-Tg mice compared to wild-type littermate controls. IOPr soma size was unchanged between genotypes (**Fig. 4C,G**), as did IO GAD immunoreactivity (**Fig. 4H,I**), revealing that inhibitory terminals on the SCA1-Tg IO are structurally intact. Based on these results, there does not appear to be a degenerative hypertrophy phenotype in the SCA1-Tg IO.

**Figure 4.**
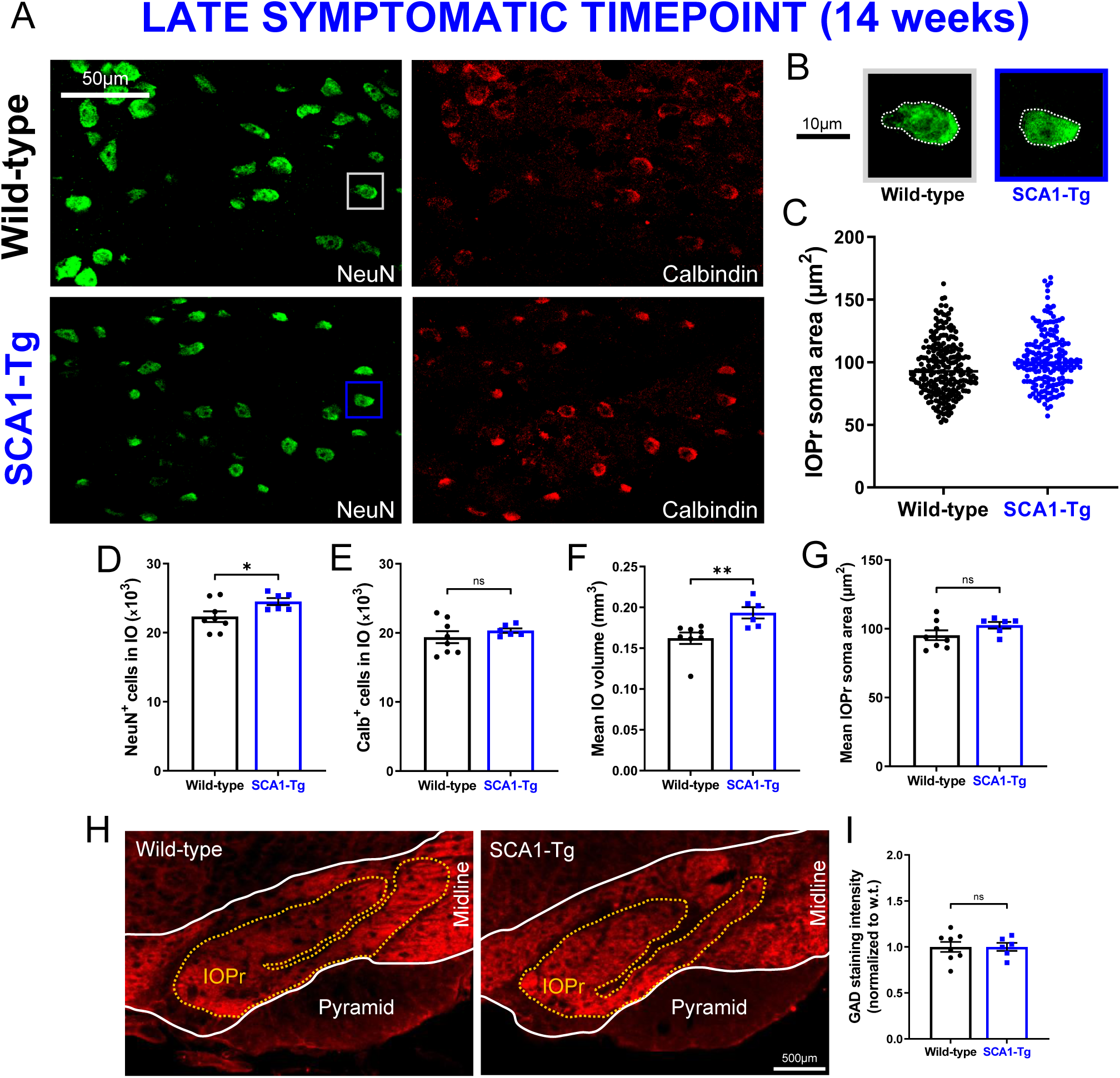
IO neuron size and viability is retained in a Purkinje cell-specific mouse model of SCA1. **(A)** Coronal histological sections showing IOPr neurons in wild-type (top) and SCA1-Tg (bottom) mice at 14 weeks. Unlike SCA1-KI mice, SCA1-Tg mice express mutant *ATXN1* solely in Purkinje cells of the cerebellum (i.e., not in the IO). These mice have a more severe phenotype, making 14 weeks of age a late symptomatic timepoint. Sections have been stained for NeuN (green) and Calbindin (red). **(B)** Inset showing representative IOPr cells of each genotype. **(C, G)** IOPr neuron soma size was estimated by measuring the area of the soma in ∼200 cells per genotype. No change in IOPr soma size in SCA1-Tg mice is evident at this timepoint. **(D-F)** Quantification of unbiased stereological analysis, providing an estimate of both total cell number and gross volume for the entire IO. At 14 weeks, degeneration is not apparent in the SCA1-Tg IO. **(H)** Coronal histological sections showing principal IO in wild-type (left) and SCA1-Tg (right) mice at 14 weeks. Sections have been stained for glutamate decarboxylase (GAD), a marker of GABAergic terminals (red). **(I)** Quantification of staining reveals that GAD signal is retained in the SCA1-Tg IO at 14 weeks, indicating no loss of inhibitory synaptic input in these mice. Data are expressed as mean ± SEM. Statistical significance derived by unpaired *t*-test with Welch’s correction, * = *P* < 0.05, ns = not significant. **(A-G)** *N_wild-type_* = 8 mice, *N_SCA1-Tg_* = 6 mice **(H,I)** *N_wild-type_* = 8 mice, *N_SCA1-Tg_* = 6 mice.

HOD has been historically understood as a non-cell-autonomous phenomenon, arising after inhibitory deafferentation of the IO (Duvoisin, 1984; Boesten and Voogd, 1985; de Zeeuw et al., 1990; Ferrer et al., 1994). These results demonstrate that, in the context of SCA1, a similar cellular phenotype can occur in the absence of any change in IO inhibitory tone. Indeed, a non-olivary disruption in the IO circuit, modeled by SCA1-Tg mice, seemed to have no effect on IO neuron size or viability. This reveals that the IO phenotype observed in SCA1-KI mice is likely cell-autonomous, suggesting a novel cause of degenerative hypertrophy separate from any extrinsic source (as in HOD).

### SCA1-KI IOPr neurons are hyperexcitable

To explore potential intrinsic causes of degenerative hypertrophy in the SCA1 IO, we performed patch-clamp electrophysiology on acute brainstem slices from 14-week-old SCA1-KI mice and wild-type littermate controls (**Fig. 5A**). All recordings were done in the IOPr, which was distinguished by its laminar structure and anatomical position lateral to the midline. The identity of targeted cells was confirmed by the presence of spontaneous subthreshold oscillations (SSTOs), a low-frequency fluctuation in membrane potential that has been well-established as a characteristic feature of IOPr neurons (Llinas and Yarom, 1986; Llinas, 2009; Choi et al., 2010) (**Fig. 5B**). Using a whole-cell current clamp configuration, we observed the spontaneous activity of these neurons for ∼100s. Due to the persistence of SSTOs throughout these recordings, resting membrane potential could not be directly calculated; however, an estimate was made by taking the average potential across 10 s of oscillations (**Fig. 5C**). This estimate of resting membrane potential, as well as the average frequency of SSTOs (**Fig. 5D**), was not significantly different between SCA1-KI IOPr neurons and wild-type IOPr neurons. SCA1-KI IOPr neurons did, however, exhibit a decreased average SSTO amplitude (**Fig. 5E**) and spontaneous spike frequency (**Fig. 5F**).

**Figure 5.**
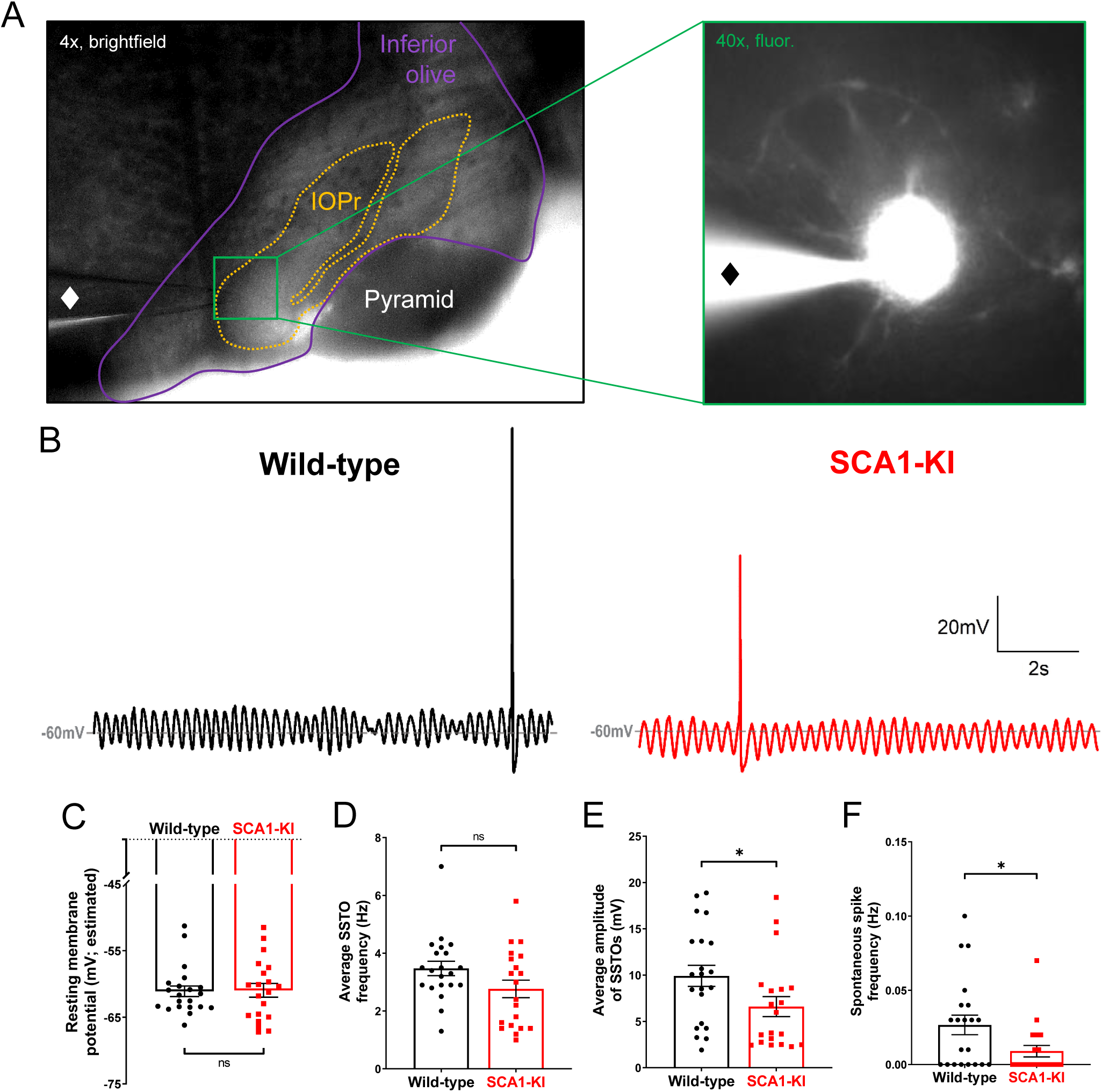
Spontaneous activity in SCA1-KI IOPr neurons is largely unchanged. **(A)** Images demonstrating whole-cell patch-clamp electrophysiology on IO neurons in coronal brainstem slices at 4x magnification (left) and 40x magnification (right). Internal solution included a small amount of fluorescent dye that filled cells while recording, allowing for the single-cell morphological analysis shown in Fig. 1. A ♦ symbol marks the recording electrode. **(B)** Representative traces of spontaneous activity in IOPr neurons from wild-type (left) and SCA1-KI (right) mice at an early symptomatic timepoint of 14 weeks. The presence of spontaneous subthreshold oscillations (SSTOs) was used as confirmation of cell type when recording. **(C-F)** Quantification of various electrophysiological properties show few differences between SCA1-KI and wild-type IOPr neurons at rest. There was no apparent change in resting membrane potential**^⸸^ (C)** or SSTO frequency **(D)** between genotypes, though SCA1-KI IOPr neurons did exhibit diminished SSTO amplitude **(E)** and spontaneous spike generation **(F)**. Data are expressed as mean ± SEM. Statistical significance derived by unpaired *t*-test with Welch’s correction, * = *P* < 0.05, ns = not significant. **^⸸^**Due to the persistence of SSTOs, the average membrane potential across 10 s or recording was used as an estimate of resting membrane potential. *N_wild-type_* = 21 cells from 15 mice, *N_SCA1-KI_* = 20 cells from 13 mice.

After recording spontaneous activity in IOPr neurons, we assessed their evoked activity by injecting increasing levels of current in a whole-cell voltage clamp configuration. Preliminary tests revealed that holding cells at a slightly hyperpolarized potential (−80 mV) limited most active conductances. This allowed for the injection of current from a stable baseline, free of SSTOs, spontaneous spikes, and other potential confounding activity. From −80 mV, we injected 0-800 pA depolarizing current in repeated 1 second steps, increasing the amount injected by +50 pA with each successive sweep (**Fig. 6B**). This generated the characteristically wide spikes previously described in IOPr neurons, with their distinctive large afterdepolarization (ADP) visible as a protruding “hump” in the repolarizing phase of the spike (Llinas and Yarom, 1981b) (**Fig. 6C**).

**Figure 6.**
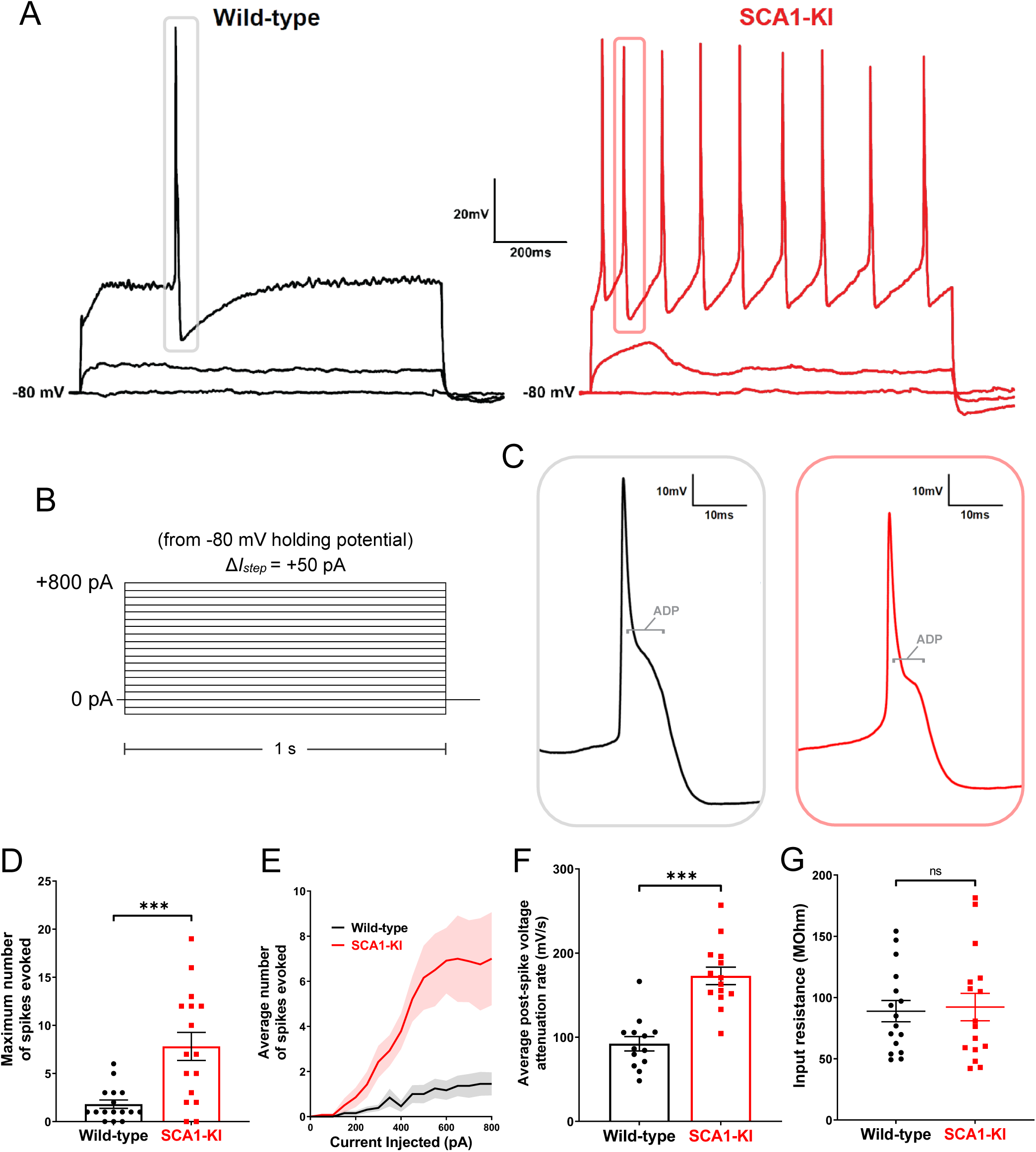
IOPr neurons in SCA1-KI mice are hyperexcitable. **(A)** Representative traces of evoked activity in IO neurons from wild-type (left) and SCA1-KI (right) mice at an early symptomatic timepoint (14wks). Traces shown for each group are 0 pA injected (bottom trace), +200 pA injected (middle trace), and +550 pA injected (top trace). **(B)** From a holding potential of −80 mV, a range of 0-800 pA depolarizing currents were injected in +50 pA, 1 second steps. Recordings were performed in current clamp mode. **(C)** Inset of representative evoked spikes at a higher timescale resolution reveal a long afterdepolarization (ADP) “hump,” a characteristic feature of IOPr neurons. **(D)** Unlike IO neurons in wild-type mice, IO neurons in SCA1-KI mice are able to sustain a spike train. **(E)** Input-output curve of average spikes produced in IO neurons of each genotype. Number of spikes rose steadily with current injection in SCA1-KI IO neurons, while wild-type IO neurons rarely exhibited spiking in the range depicted (+0-800 pA injected). **(F)** The rate at which membrane potential recovered back to baseline from its minimum value post-spike was significantly higher in SCA1-KI IOPr neurons, allowing these cells to fire repetitively. **(G)** Input resistance of IO neurons is unchanged in SCA1-KI mice, demonstrating that this hyperexcitability phenotype is not a product of any change in voltage generated per injected current step. Data are expressed as mean ± SEM. Statistical significance derived by unpaired *t*-test with Welch’s correction, *** = *P* < 0.001, ns = not significant. **(D,E,G)** *N_wild-type_* = 16 cells from 14 mice, *N_SCA1-KI_* = 16 cells from 11 mice **(F)** Non-firing cells removed, so *N_wild-type_* = 13 cells from 12 mice, *N_SCA1-KI_* = 14 cells from 9 mice.

Current injections in the IOPr revealed a novel hyperexcitability phenotype in the SCA1 IO (**Fig. 6A**). Unlike wild-type IOPr neurons, SCA1-KI IOPr neurons are able to fire repetitively (**Fig. 6D**) in spike trains that increase in frequency with greater injections of current (**Fig. 6E**). Input resistance was unchanged between the two genotypes (**Fig. 6G**), indicating that differences in excitability could not be explained by changes in passive membrane properties. Rather, further analysis of these spikes suggests that the observed SCA1-KI IOPr hyperexcitability may be related to a disruption of currents post-spike. In SCA1-KI mice, the rate at which IOPr membrane potential depolarizes back to baseline from its minimum is nearly two-fold the rate observed in wild-type mice (**Fig. 6F**). This suggests that the membrane potential of SCA1-KI IOPr neurons can more quickly recover from a previous spike and reach threshold again, which may explain how these cells, unlike wild-type IOPr neurons, are able to generate spike trains.

### Ion channel transcripts are dysregulated in the SCA1-KI medulla

In order to assess potential causes of hyperexcitability in the SCA1-KI IO, we analyzed previously-obtained medullary transcriptome data from SCA1-KI mice treated with antisense oligo-nucleotides (ASOs) to block *Atxn1* expression. After being administered ASOs at 5 weeks of age, these mice demonstrated significant rescue in both motor behavior and brainstem phenotypes. (Friedrich et al., 2018). At 28 weeks, medullary tissue was harvested from both treated and non-treated mice, as well as untreated wild-type littermate controls, and analyzed by RNA-seq. Comparing untreated wild-type and untreated SCA1-KI medullary transcripts revealed significant dysregulation in 1374 genes. Interestingly, 31 ion channel transcripts were present among these baseline differentially-expressed genes (DEGs), representing a 2.79-fold enrichment compared to ion channel representation in the mouse genome. This enrichment is considered significant (Fisher’s exact test, *P* = 1.41_E_-6), suggesting that disrupted ion channel expression could play a key role in the development of SCA1-KI brainstem phenotypes. Of these 31 candidate genes, 6 encode ion channels that influence excitability synaptically (IC_synaptic_), while the remaining 25 encode ion channels that influence excitability intrinsically (IC_intrinsic_) (Alexander et al., 2011). For all but one of these channels, medullary transcripts were downregulated in SCA1-KI mice compared to wild-type controls (**Fig. 7A**), a finding that comports with previous studies that report a primarily downregulated DEG population in SCA1 mouse models (Lin et al., 2000; Niewiadomska-Cimicka et al., 2020).

**Figure 7.**
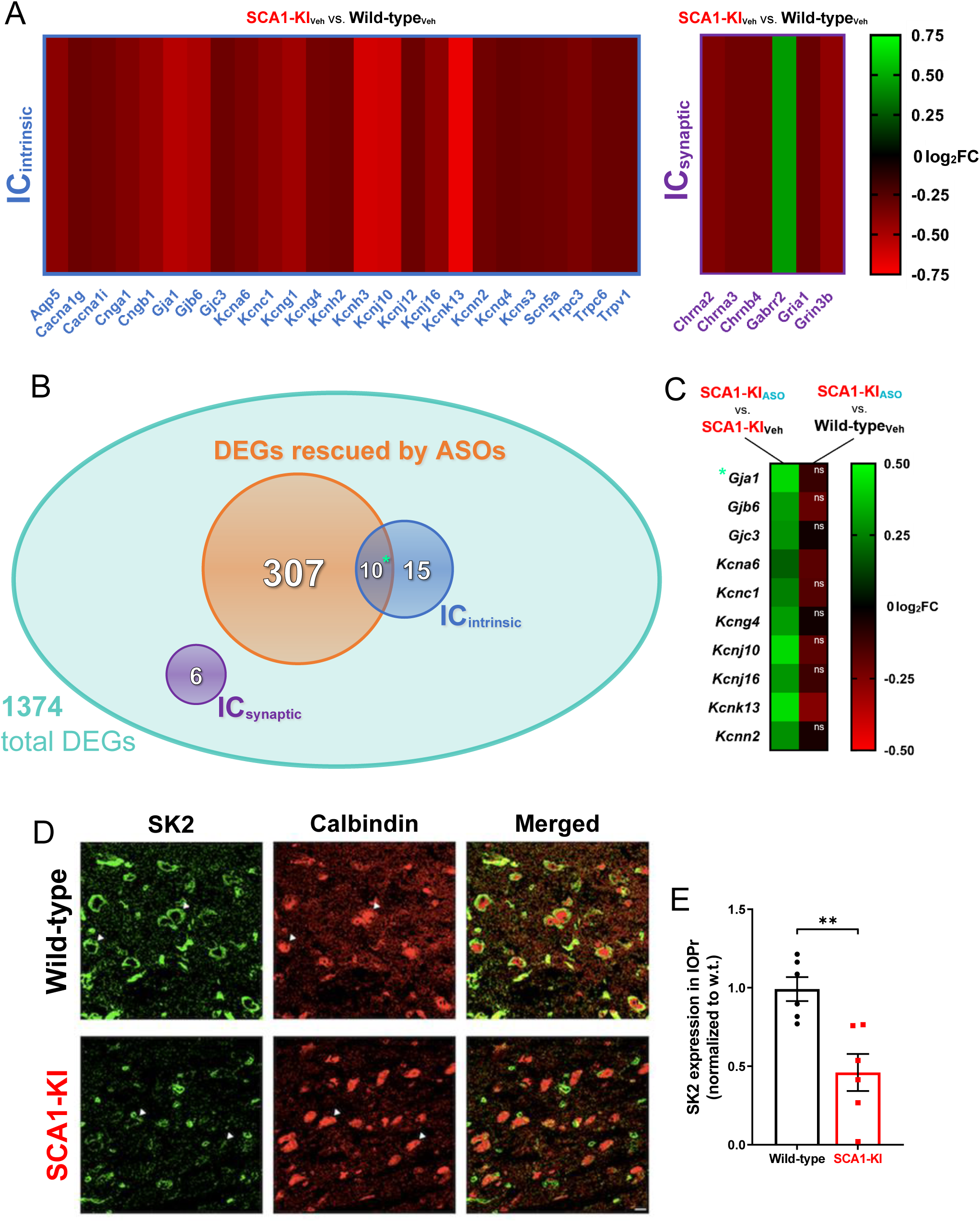
Ion channel expression is disrupted in the SCA1-KI medulla. **(A)** Transcriptomic analysis of whole medulla from SCA1-KI and wild-type mice at 28 weeks reveal that 31 ion channel genes are differentially expressed. Of these, 25 influence excitability intrinsically (IC_intrinsic_), while 6 influence excitability synaptically (IC_synaptic_). **(B)** Downregulation of *Atxn1* expression by antisense oligonucleotides (ASOs) ameliorates brainstem phenotypes in SCA1-KI mice. Of the 1374 differentially expressed genes (DEGs) in the SCA1-KI medulla, 317 DEGs were significantly rescued by ASO treatment (and, therefore, are more likely to be responsible for the observed phenotypic rescue). Within this cohort, IC_intrinsic_ genes were significantly enriched. **(C)** Heatmap showing differential expression of the 10 IC_intrinsic_ DEGs between treatment groups. These ion channel genes exhibited a significant increase in expression after ASO treatment (left), with the majority of them rising to wild-type levels (right). **(D)** In order to examine how decreases in medullary ion channel transcripts may result in a loss of channels in the IOPr, immunostaining for the small-conductance potassium channel SK2 was performed. Coronal histological sections of the IOPr in wild-type (top) and SCA1-KI (bottom) mice at 14 weeks (an early symptomatic timepoint) are shown. Sections have been stained for SK2 (green) and calbindin (red). **(E)** Quantification of immunostaining reveals a significant loss of SK2 channels on the membrane of SCA1-KI IOPr neurons. Data are expressed as mean ± SEM. Statistical significance derived by Fischer’s exact test (enrichment analyses) or unpaired *t*-test with Welch’s correction (all other comparisons), ** = *P* < 0.01, ns = not significant. **(A-C)** *N_wild-type, Vehicle_* = 8 mice, *N_SCA1-KI, Vehicle_* = 8 mice, *N_SCA1-KI, ASO_* = 7 mice. **(D, E)** *N_wild-type_* = 6 mice, *N_SCA1-KI_* = 6 mice.

To further narrow down this list of potential candidates genes, we analyzed data amongst the ASO treatment groups. Of the original 1374 medullary DEGs in SCA1-KI mice, 317 showed significant rescue after ASO treatment. Included among these 317 ASO-rescued DEGs were 10 ion channel genes, all of which encoded IC_intrinsic_ channels (**Fig. 7B**). This constituted a significant 5.26-fold enrichment of IC_intrinsic_ genes compared to IC_intrinsic_ representation in the mouse genome (Fisher’s exact test, *P* = 3.43_E_-5), suggesting that the loss of ion channels that regulate intrinsic excitability may be an important contributing factor to brainstem dysfunction in SCA1-KI mice. Of the IC_intrinsic_ candidate genes that underwent significant rescue by ASO treatment, the majority (8 of the 10) achieved “complete” rescue; i.e., their expression level after ASO treatment was not significantly different from baseline levels in wild-type mice (**Fig. 7C**). The full medullary transcriptome dataset analyzed here is available in **Table 7-1**, which reports raw reads for all 30,973 transcripts assessed in each group, as well as the results of the statistical comparisons between groups used to define DEGs.

To determine whether this decrease in medullary IC_intrinsic_ transcripts reflects a loss of protein levels in the IO, we immunostained coronal brainstem slices from 30-week-old SCA1-KI and wild-type mice for the channel encoded by *Kcnn2*: small conductance calcium-activated potassium channel 2 (SK2) (**Fig. 7D**). Quantification of SK2 staining intensity on IOPr neurons reveals a ∼50% reduction in protein levels (**Fig. 7E**). Taken together, these results suggest that downregulation of IC_intrinsic_ genes in the medulla may play a critical role in SCA1 brainstem pathogenesis.

### Spikes from SCA1-KI IOPr neurons exhibit a diminished AHP

In order to assess the functional consequence of IC_intrinsic_ channel loss in the medulla, we recorded from IOPr neurons using patch-clamp electrophysiology in acute brainstem slices. Using the same voltage clamp protocol described above (**Fig. 8A**), increasing levels of current were injected from a holding potential of −80 mV in 1 s steps to generate spikes. AHP depth from each spike generated from 0 to 800 pA current injected was normalized to that spike’s threshold and compiled (note: two hyperpolarizing currents, −100 pA and −50 pA, were injected at the beginning of this protocol to allow for input resistance calculations before interference by spike generation). This revealed a significant loss of average AHP depth in SCA1-KI IOPr neurons (**Fig. 8B,C**). To determine whether this phenomenon is also capable of occurring in IOPr neurons at rest, we conducted similar spike analysis experiments, this time generating single spikes from −60 mV (a holding potential close to the estimated resting membrane potential of these cells (**Fig. 5C**)). In the whole-cell voltage clamp configuration, we injected 0-1000 pA depolarizing current in repeated 10 ms steps, increasing the amount injected by +50 pA with each successive sweep (**Fig. 8D**). In this setting, without the presence of spike trains, average AHP depth was again diminished in SCA1-KI IOPr neurons compared to wild-type IOPr neurons (**Fig. 8F**). Previous studies have demonstrated that potassium channels are the primary determinants of AHP size and shape in IOPr neurons. The observed AHP deficit in SCA1-KI IOPr neurons, as well as the enrichment of IC_intrinsic_ genes among ASO-rescued DEGs in the SCA1-KI medulla (7 of which encode potassium channels (**Fig. 7C**)), connects ion channel loss in the SCA1 brainstem to a potential functional consequence in IOPr hyperexcitability.

**Figure 8.**
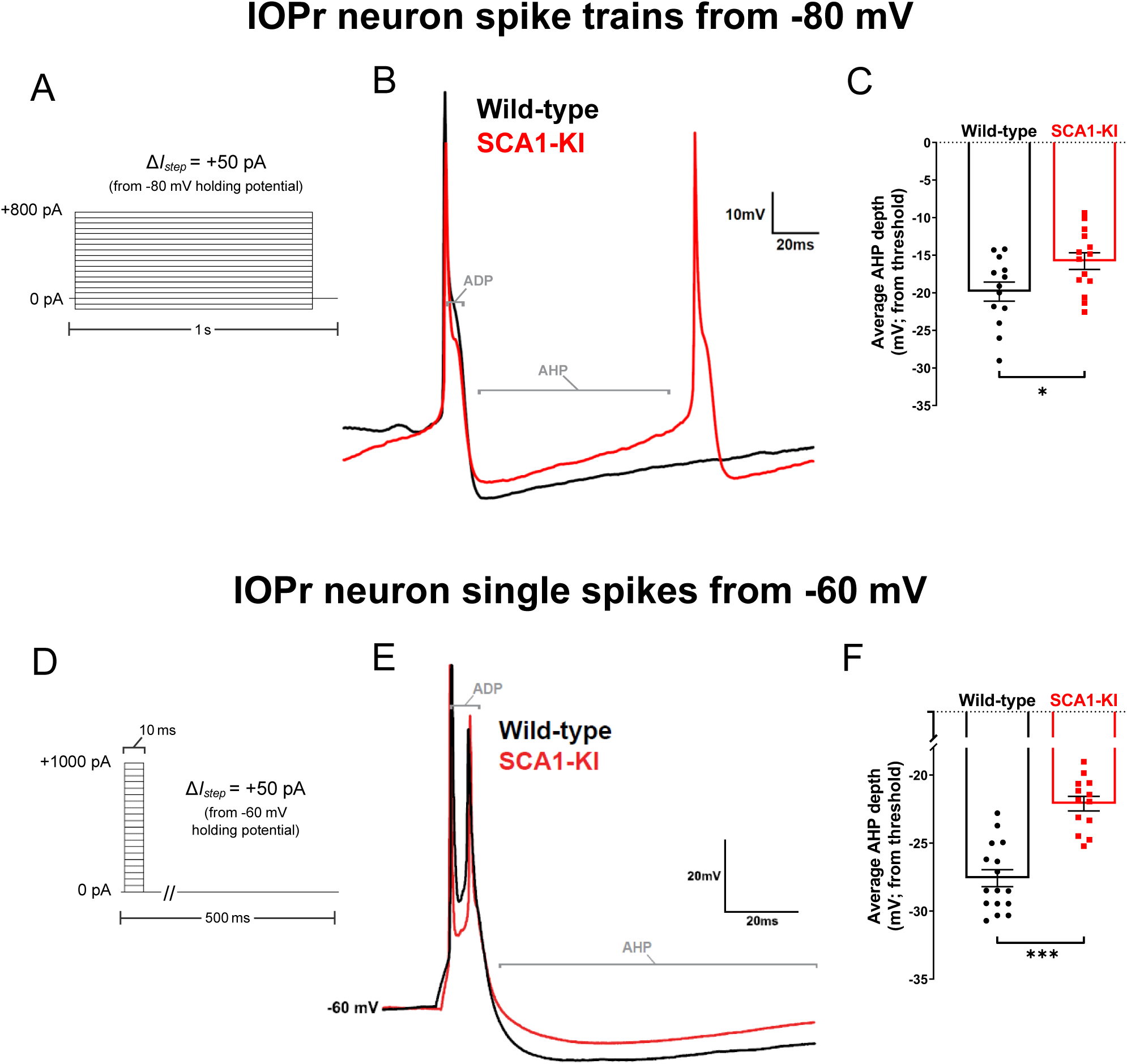
Spike afterhyperpolarization is diminished in SCA1-KI IOPr neurons. **(A)** In order to assess the functional relevance of ion channel loss in SCA1-KI IOPr neurons, spike trains were generated by injecting increasing levels of current from −80 mV (a holding potential at which active conductances are minimal in the IO). **(B)** Representative traces of spikes generated by injecting +450 pA current in IOPr neurons of wild-type and SCA1-KI mice at 14 weeks. **(C)** Quantification of AHP depth (measured from each spike’s threshold), demonstrating that SCA1-KI IOPr spikes during spike trains exhibit a shallower AHP compared to wild-type. **(D)** In order to determine if this AHP loss can also occur at rest, single spikes were generated by injecting increasing levels of current from −60 mV (a holding potential close to resting V_m_). Because the amount of current needed to generate a single spike varied between cells, the first spike that exhibited an ADP was used for comparisons. **(E)** Representative traces of evoked activity in IOPr neurons in wild-type and SCA1-KI mice at 14 weeks are overlaid. **(F)** Quantification of AHP depth (measured from each spike’s threshold), demonstrating that SCA1-KI IO neuron spikes from rest also exhibit a shallower AHP compared to wild-type. Data are expressed as mean ± SEM. Statistical significance derived by unpaired *t*-test with Welch’s correction, * = *P* < 0.05, *** = *P* < 0.001. **(A-C)** *N_wild-type_* = 13 cells from 12 mice, *N_SCA1-KI_* = 14 cells from 9 mice. **(D-F)** *N_wild-type_* = 16 cells from 14 mice, *N_SCA1-KI_* = 13 cells from 9 mice.

## Discussion

Though the IO has long been identified as a characteristic area of SCA1 pathology (Seidel et al., 2012), it is not known what mechanisms drive disease in this cell population. Here, we describe degenerative hypertrophy in SCA1 IOPr neurons, a previously-unknown IO phenotype of the SCA1-KI mouse model. This appears strikingly similar to hypertrophic olivary degeneration (HOD), a pathology caused by the loss of inhibitory afferents to the IO (Ruigrok et al., 1990; Wang et al., 2019). Our results, however, demonstrate that inhibitory innervation to the IO in SCA1-KI mice is both structurally and functionally intact. Though some limited occurrence of HOD in the context of SCA has been previously reported (Yoshii et al., 2017), this study constitutes the first description of a robust HOD-like phenotype in the proven absence of inhibitory deafferentation. This suggests that there is some downstream consequence of synaptic disinhibition, rather than synaptic disinhibition itself, that is both necessary and sufficient to cause HOD.

HOD, as historically described, is caused by a lesion in the central tegmental tract or superior cerebellar peduncle, consistent with loss of IO innervation from the dentate nucleus of the cerebellum (Sabat et al., 2016; Tilikete and Desestret, 2017). Though IO neurons receive innervation from a variety of brain regions, previous studies have shown that the dentate nucleus supplies the IO with its primary inhibitory input (Fredette and Mugnaini, 1991; Best and Regehr, 2009). Importantly, disruption of this dentato-olivary circuit is the sole condition in which HOD has been observed, leading to the current conclusion that HOD is a consequence of IO neurons losing inhibitory tone. Though we found inhibitory afferents to be intact in the SCA1 IO, these neurons still exhibit hyperexcitability, indicating that increased membrane excitability may be HOD’s proximal cause; that is, HOD may occur as the result of *any* phenomenon that sufficiently increases IO membrane excitability, be it extrinsic (as in HOD, historically) or intrinsic (as in SCA1).

One potential explanation for this connection between excitability and cell size is the possibility that, in the SCA1 IO, hypertrophy is acting in opposition to hyperexcitability as a compensatory mechanism. A substantial lengthening of dendrites, as exhibited in SCA1-KI mice at 14 weeks (**Fig. 1**), is likely to enhance dendritic shunting (Blomfield, 1974). If the multiple excitatory signals that the IO receives were sufficiently diluted by shunting through this extra length of dendrite, membrane excitability would, effectively, be decreased. This may explain the reduction in SCA1-KI Purkinje cell complex spike frequency we observed *in-vivo*: i.e., even though synaptic input to the SCA1-KI IO appears intact, shunting could be diminishing the number of excitatory signals that end up reaching the soma and triggering IO output to the cerebellum – a result that would likely cause deficits in motor learning (Lang et al., 2017). This raises the interesting possibility that HOD may be a ‘rogue’ compensatory mechanism, such that IO hypertrophy helps the cell (by alleviating hyperexcitability) but harms the circuit (by reducing final cellular output). Similarly, previous studies provide evidence that changes in morphology may also be acting as a compensatory mechanism against hyperexcitability in SCA1 Purkinje cells. Prior to atrophy, Purkinje cells in SCA1-Tg mice exhibit a severe firing phenotype caused by increased membrane excitability; however, after the onset of atrophy, there is a partial restoration of firing. This is likely due to changes in channel density on SCA1 Purkinje cell somas. SCA1 Purkinje cells exhibit reduced expression of multiple key ion channel genes, causing a significant decrease in somatic channel density. However, as a result of a decrease in somatic surface area during atrophy, channel density on the soma increases, which appears to partially alleviate the SCA1-Tg Purkinje cell firing phenotype by reducing membrane excitability (Dell’Orco et al., 2015). Though it is unclear how this compensatory effect of atrophy occurs, there is evidence that channel density on the Purkinje cell soma is an important factor in the formation of functional ion channel clusters, especially those that regulate membrane excitability via potassium conductances (Womack et al., 2004; Kaufmann et al., 2009; Indriati et al., 2013). Taken together, these results suggest a close relationship between morphology and excitability in neurons that are selectively vulnerable to SCA1-associated degeneration. In addition, it raises the question of what causal link, if any, exists between the structural and functional pathologies of these neurons.

Previous research has demonstrated that the mutation that causes SCA1 (an expansion in the polyglutamine-encoding CAG repeat region of the *ATXN1* gene) produces Purkinje cell pathology by disrupting the expression of a host of downstream genes (Lin et al., 2000; Crespo-Barreto et al., 2010). Due to an abnormal interaction between the polyglutamine-expanded ATXN1 protein and Capicua (CIC), a transcriptional repressor, the vast majority of these Purkinje cell genes exhibit decreased expression in SCA1 (Lam et al., 2006; Rousseaux et al., 2018). Ion channel genes, including many that are crucial for maintaining Purkinje cell excitability, are significantly enriched among this group (Bushart et al., 2018; Chopra et al., 2020). Similarly, in this study, we found a significant enrichment of ion channel genes among the medullary DEGs identified in 28-week-old SCA1-KI mice. the majority of which also exhibit decreased expression. Interestingly, cerebellar transcripts from these same 28-week-old SCA1-KI mice reveal that DEGs in the SCA1 cerebellum and the SCA1 medulla are indeed similar, but not identical (Friedrich et al., 2018). Between these two tissues, 7 ion channel DEGs are shared: *Cacna1g*, *Gria1*, *Kcna6*, *Kcnc1*, *Kcnk13*, *Kcnn2*, and *Trpc3*, all exhibiting decreased expression in both brain regions. Based on these observations, it appears that the cerebellum and brainstem may share a common mechanism of SCA1 pathology in reduced ion channel expression. This connection might be explained by the shared presence of CIC, which is moderately expressed in both the mouse cerebellum and the mouse IO (Lein et al., 2007). The discrepancies between 28 week SCA1-KI medullary and cerebellar DEGs, as suggested by a recent study (Driessen et al., 2018), may be due to the involvement of transcription factors other than CIC that are also affected by polyglutamine-expanded ATXN1.

Within the SCA1 brainstem and cerebellum, there appears to be additional convergence that links IO neuron and Purkinje cell pathology, specifically. Previous research has shown that loss of channels in SCA1-KI Purkinje cells causes a firing dysfunction related to an increase in membrane excitability (Dell’Orco et al., 2015; Chopra et al., 2018). This is driven primarily by abnormalities in spike afterhyperpolarization (AHP), which is significantly shallower in SCA1-KI Purkinje cells at 14 weeks (Bushart et al., 2021). SCA1-KI IO neurons, shown by their ability to generate spike trains (**Fig. 6**), also exhibit an increase in membrane excitability that correlates with an observed AHP deficit (**Fig. 8**). This suggests that hyperexcitability caused by a reduction in spike AHP is a phenotype that is shared by both IO neurons and Purkinje cells in SCA1-KI mice.

Despite differences in their baseline firing properties, AHP dynamics in both Purkinje cells (Edgerton and Reinhart, 2003; Walter et al., 2006) and IO neurons (Llinas and Yarom, 1981b; Lang et al., 1997) appear to be largely governed by the interplay between calcium currents and potassium currents – especially via the conductance of calcium-activated potassium (K_Ca_) channels. Of the channels that are represented in the two groups of 28 week SCA1-KI DEGs, the majority are either calcium or potassium channels. In addition, one of the 7 channel DEGs shared by both the SCA1-KI cerebellar and medullary datasets is *Kcnn2*, which encodes one of only 5 K_Ca_ channels expressed in the brain (Alexander et al., 2017). Together, these findings suggest that hyperexcitability due to altered calcium and/or potassium conductances may represent a shared mechanism of dysfunction in SCA1 IO neurons and Purkinje cells.

Though the relationship between these physiological changes and cell size remains unclear, prior research has demonstrated that firing abnormalities in SCA1-KI Purkinje cells precede degeneration (Hourez et al., 2011). Previous work has shown that addressing this by increasing specific potassium conductances with pharmacological agents can faithfully restore SCA1-KI Purkinje cell firing *in-vitro*. Additionally, chronic treatment of SCA1-KI mice with these same drugs not only improves motor function but, importantly, also slows the atrophy and subsequent degeneration of Purkinje cells (Chopra et al., 2018; Bushart et al., 2021). As such, by demonstrating that a reduction in SCA1-KI Purkinje cell hyperexcitability is sufficient to partially rescue cell size phenotypes, these results suggest that SCA1-associated disruptions of membrane excitability may be an important driver of morphological changes in Purkinje cells. Similarly, though it is not known what specific effect it had on either SCA1-KI IO neuron hyperexcitability or hypertrophy, ASO administration did partially alleviate brainstem phenotypes in these mice (Friedrich et al., 2018). In addition, the enrichment of ion channel genes among the SCA1-KI medullary DEGs rescued by ASOs (**Fig. 7**) suggests that membrane excitability might be one of the cellular features whose restoration is important to this ASO-mediated rescue of brainstem dysfunction. Together, these results indicate that altered membrane excitability may constitute a key driver of SCA1 pathology in the brainstem – including HOD.

## Conflicts of Interests

The authors declare no competing financial interests.

## Acknowledgments

This work was supported by funding from the US National Institutes of Health/National Institute of Neurological Disorders and Stroke (NIH/NINDS) (R01 NS085054) and a National Ataxia Foundation Research Seed Money Grant Award to VGS, as well as funding from the NIH/NINDS (R37 NS022920), a National Ataxia Foundation Pioneer Award, and a Wallin Neuroscience Discovery Award to HTO.

## Notes

### Competing Interest Statement

The authors have declared no competing interest.

